# Estimating self-fertilization rates from the diploid genotype spectrum

**DOI:** 10.64898/2026.10.13.751067

**Authors:** Patrick F. McKenzie, Felix L. Wu, Robin Hopkins

## Abstract

The distribution of self-fertilization across the tree of life remains poorly characterized. Here we present the diploid genotype spectrum (DGS) as a useful genomic data summary for characterizing populations undergoing partial self-fertilization (selfing). This statistic provides a straightforward baseline for estimating selfing rates from population genomic data. At every segregating site across a genome, the diploid genotype configuration records the numbers of homozygous ancestral, heterozygous, and homozygous derived individuals, and the DGS compiles the counts of these configurations across sites. We show that the expected relative frequencies of DGS configurations among segregating sites can be computed under the standard coalescent approximation for partial selfing. We present two methods for doing this, one that is an intuitive recursion through possible states of the model, and one that is a more efficient combinatorial version. We give a worked example, validate the analytical expectations using forward simulations, and introduce a software package that estimates population selfing rates from diploid SNP data. The computations here mechanistically link selfing to observed patterns of diploid genotype configurations.

## 1 Introduction

Self-fertilization (selfing) has evolved repeatedly among hermaphroditic organisms, despite substantial costs. By providing reproductive assurance under mate limitation, selfing can increase the probability of successful reproduction, but it also decreases heterozygosity and can expose recessive deleterious variation (Shimizu and Tsuchimatsu, 2015). Understanding the circumstances that favor the evolution of selfing requires knowing how frequently selfing occurs across taxa. However, the distribution of selfing rates across the tree of life remains poorly understood, in part because measuring selfing rates has historically depended on a combination of intensive natural history observations and a limited set of genetic methods (Goodwillie et al., 2005; Jarne and Auld, 2006; Meyer et al., 2025, 2026; Shimizu and Tsuchimatsu, 2015; Soltis and Soltis, 1992). Improving our understanding of the frequency of selfing requires genome summaries that capture its effects on genetic variation.

To see the kind of information that such a summary might use, consider a segregating site at which the derived allele is present in two copies in a sample of diploid individuals. Those copies could occur together in one homozygous-derived individual, or instead perhaps separately in two heterozygous individuals. The ordinary site frequency spectrum (SFS) only records the total number of derived alleles (Gutenkunst et al., 2009; Watterson, 1984), so it is unable to distinguish between these two possible genotype configurations. Under selfing, though, these two configurations are usually not equally probable. Selfing increases the probability that the two homologs within an individual share recent ancestry (Nordborg and Donnelly, 1997; Shimizu and Tsuchimatsu, 2015) and so alters how derived alleles are distributed among diploid genotypes.

Ferretti et al. (2018) formalized this additional single-site information with the joint site frequency-heterozygosity spectrum *ψ*(*f, h*), which classifies segregating sites jointly by derived-allele frequency *f* and the fraction of sampled individuals that are heterozygous, *h*. These values completely specify the numbers of ancestral homozygotes, heterozygotes, and derived homozygotes at a biallelic site. Here we use an equivalent reparameterization that we call the diploid genotype spectrum (DGS). At each segregating site we record the configuration (*n*_0_, *n*_1_, *n*_2_): the numbers of individuals that are homozygous ancestral, heterozygous, and homozygous derived, respectively. The DGS records the number of sites falling into each such configuration across the genome.

Selfing changes the relative frequencies of these configurations, which can be calculated under the partial-selfing coalescent of Nordborg and Donnelly (1997). We use these expected frequencies to infer the population selfing rate from an observed DGS.

Existing genetic approaches estimate selfing from heterozygote deficiency, often summarized by inbreeding coefficients like *F*_*IS*_ (Bürkli et al., 2017; Weir and Cockerham, 1984), from correla-tions in heterozygosity among unlinked loci (identity disequilibrium) (David et al., 2007), or from genome-wide recombination and local coalescence (Strütt et al., 2023). Other approaches use the partial-selfing coalescent to relate DNA sequence variation to selfing. These include estimators based on the number of segregating sites (Fu, 1997) and moment and likelihood approaches for estimating selfing together with a scaled mutation parameter (Nordborg and Donnelly, 1997). Particularly relevant here, Redelings et al. (2015) introduced a Bayesian method that uses genotypes at multiple unlinked loci to estimate selfing, mutation parameters, and the number of consecutive selfing generations in each sampled individual’s recent ancestry. The DGS likewise retains diploid genotype information, but summarizes that information differently, pooling sites according to the numbers of ancestral homozygotes, heterozygotes, and derived homozygotes. As we show below, the expected relative genotyupe configuration frequencies among segregating sites can be calculated under the partial-selfing coalescent without estimating a separate inbreeding history for each individual. Blischak et al. (2020) used a similarly compact set of counts to study inbreeding, but that method sums over the possible diploid genotype configurations compatible with each derived allele count, yielding the more compact SFS, rather than treating the genotype configurations themselves as the observed genomic summary.

Here we derive DGS configuration probabilities directly under the standard partial-selfing coalescent approximation (Möhle, 1998; Nordborg and Donnelly, 1997), allowing rapid estimation of population selfing rates from samples containing modest numbers of individuals. We present both an explicit recursive derivation and a more efficient combinatorial calculation, we introduce a software package for constructing and analyzing the DGS, and we compare the resulting analytical expectations with forward simulations.

## 2 Diploid genotype configurations

First we will establish the foundational data structure to calculate the DGS. Consider a sample of *n* diploid individuals. At a polarized biallelic site, let *A* denote the ancestral allele and *a* denote the derived allele. Each sampled individual has one of three genotypes

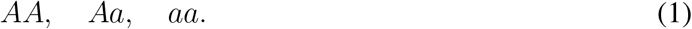

Looking across the sampled individuals, define the diploid genotype configuration of the site

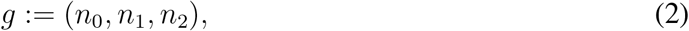

where *n*_0_ is the number of homozygous ancestral individuals, *n*_1_ is the number of heterozygous individuals, and *n*_2_ is the number of homozygous derived individuals. These counts satisfy

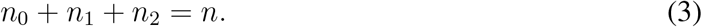

The derived allele count associated with configuration *g* is

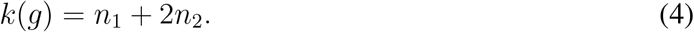

## 3 The diploid genotype spectrum

Let *G* be the set of possible polymorphic (i.e., excluding monomorphic ancestral or monomorphic derived sites) diploid genotype configurations,

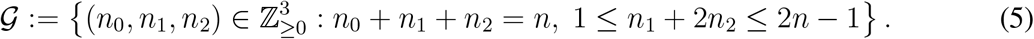

There are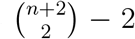of these possible polymorphic configurations (i.e.,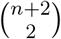 is the number of possible ways to divide *n* individuals into three groups, minus the two monomorphic cases).

For each segregating site *b*, let *g*_*b*_ ∈ *G* be its observed configuration. The DGS is the count vector

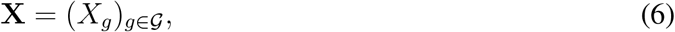

where

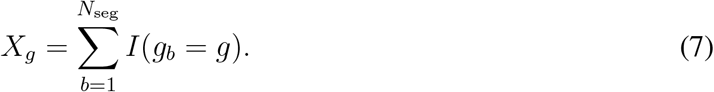

Here, *N*_seg_ is the number of segregating sites, and *I*(·) is the indicator function, returning a value of 1 when it is satisfied and 0 otherwise. So, our core object **X** just records how many sites have each possible genotype configuration.

The configurations in **X** can be grouped by their derived allele count *k* = *n*_1_ +2*n*_2_. As a simple example, for *n* = 3 sampled diploid individuals:

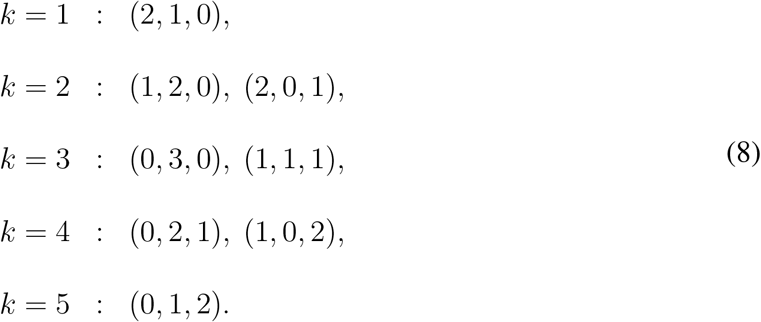

As we see here, multiple DGS configurations can map to the same SFS bin.

To formally link the DGS to the SFS, let

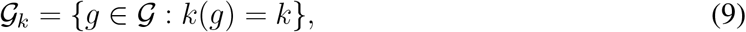

essentially grouping together DGS configurations with the same derived allele count. The SFS entry for derived allele count *k* is

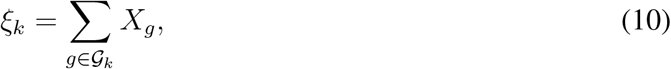

which sums up each of these same-derived-allele-count groups. Or, equivalently, the SFS is just the projection

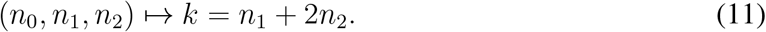

## 4 Partial selfing and the “fast”-stage collapse

We assume a neutral, panmictic population of constant size with a constant selfing rate, at equilibrium under the partial-selfing coalescent approximation.

Under the standard partial-selfing coalescent approximation, the two homologs sampled from the same diploid individual first pass backward in time through a “fast” within-individual phase during which selfing might have occurred (Möhle, 1998; Nordborg and Donnelly, 1997) (Figure 1). Here, coalescence at a locus means that the two lineages trace back to the same ancestral gene copy, not merely to the same parent. For example, ignoring mutation at that locus, an *aa* offspring of a selfed *Aa* parent inherits both copies from the parent’s *a*-bearing homolog, so the lineages coalesce in that parental generation. An *Aa* offspring instead inherits one copy from each parental homolog, so its two lineages remain distinct in that generation. If the selfing rate is *s*, the probability that the two homologs coalesce during the fast phase is

**Fig 1.**
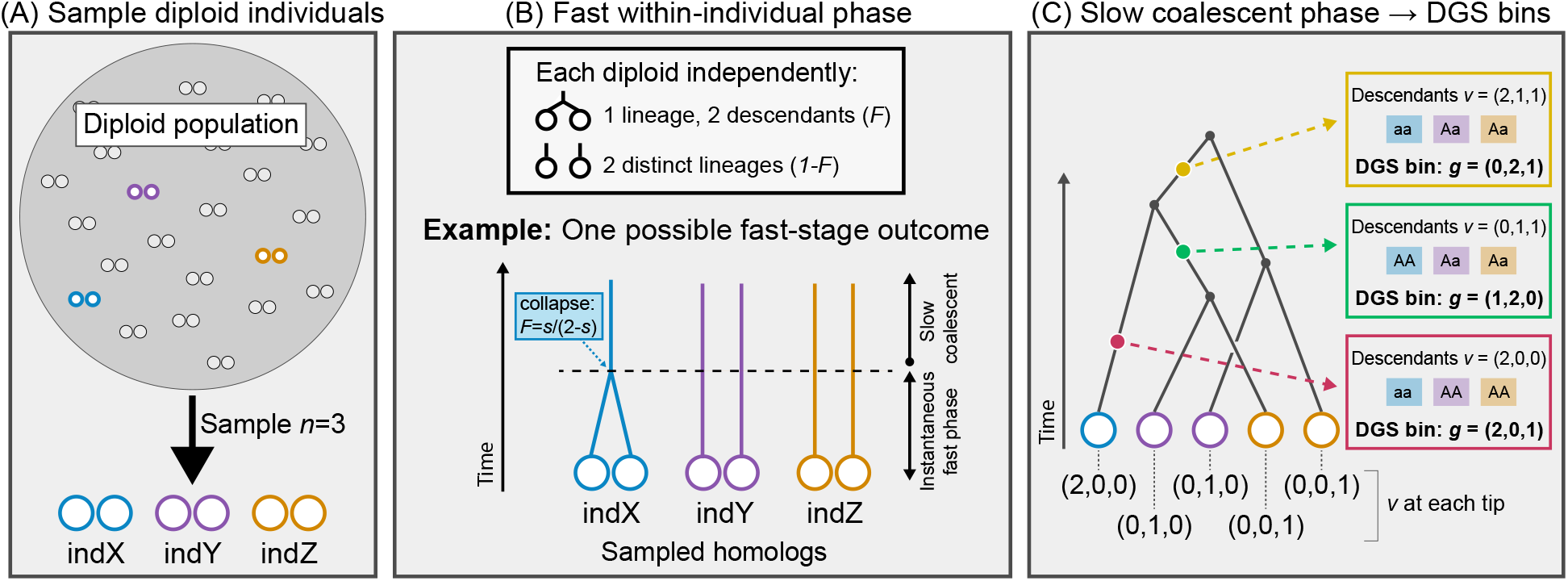
Illustration of the partial-selfing model and its relationship to the DGS. (A) Individuals in a population are represented by two touching circles corresponding to two alleles. *n* diploid individuals are sampled from a Wright-Fisher population as indicated with the colored circles. (B) The fast phase is treated as instantaneous on the population-level coalescent timescale, with its contribution to mutational edge length neglected. During this phase, the homologs of each diploid individual might collapse to form one lineage with probability 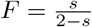, or persist into the slow phase as distinct lineages with probability 1 − *F*. (C) The slow phase consists of a backward-in-time Kingman coalescent in which remaining lineages coalesce at random. A possible coalescent tree is shown here as an example. Example mutations, each of which would generate a segregating site, are displayed on the coalescent tree as colored dots, and their corresponding descendant genotypes and DGS bin contributions are shown.

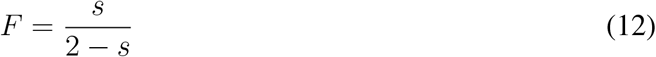

(Equation 4 of Nordborg and Donnelly (1997)). If the homologs do not coalesce, they escape into the slower population-level coalescent as two distinct lineages (details in the next section).

It is straightforward to compute the probability of different fast-stage outcomes. For a sample of *n* diploid individuals, let

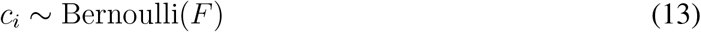

indicate whether the two homologs from individual *i* collapse during the fast phase (with 1 indicating collapse). So a fast-stage outcome is a vector

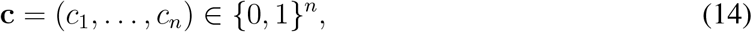

with probability

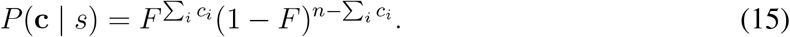

After the fast phase, the number of remaining lineages is

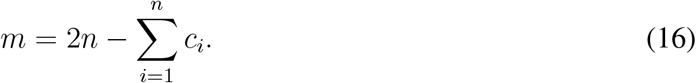

Collapsed lineages represent both homologs of a sampled individual, while uncollapsed lineages instead keep homologs separate.

Under the model of Nordborg and Donnelly (1997), the fast phase determines the initial state of the slow coalescent. It is treated as instantaneous on the population-level coalescent timescale, so its contribution to mutational edge length is neglected. It affects DGS probabilities by changing which slow lineages represent one versus two sampled homologs from an individual, whereas mutations are only assigned to edges of the genealogy during the following “slow” phase.

## 5 Expected DGS probabilities from “slow”-phase edge lengths

Under the infinite-sites model, mutations occur at a constant rate along genealogical edges. Reflecting this, the expected number of mutations producing configuration *g* is proportional to the expected total length of the edges on which a mutation would produce *g*.

With this in mind, let *G* denote the configuration of a randomly selected segregating site, and let *L*_*g*_(*s*) denote the total genealogical edge length on which a mutation would produce configuration *g*. Then, the probability of an arbitrary segregating site having configuration *g* is

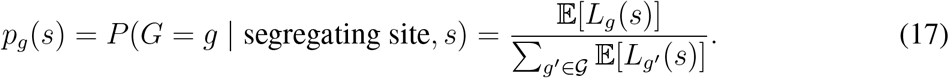

Or, the probability of observing a particular DGS configuration at the segregating site is the expected edge length on which a mutation would produce that configuration, normalized by the total expected edge length of the coalescent tree. To briefly see why, suppose the mutation rate is a constant *µ*. The expected number of mutations producing configuration *g* is then proportional to

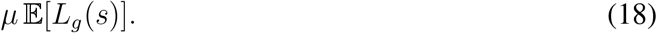

The mutation rate multiplies the expected counts for all of the configurations equally, so it cancels out when those counts are converted to relative probabilities, as is the case when we are conditioning on observing a segregating site. A result of this is that *p*_*g*_(*s*) describes the relative frequencies of DGS configurations among segregating sites rather than predicting the total number of segregating sites.

During each interval between coalescences, every extant lineage creates a genealogical edge. We can label that edge by the sampled homologs that descend from it

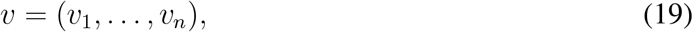

where *v*_*i*_ ∈ {0, 1, 2} is the number of sampled homologs from diploid individual *i* that descend from that edge (illustrated in Figure 1). A mutation on an edge with the descendant vector *v* is absent from individual *i* if *v*_*i*_ = 0, heterozygous in individual *i* if *v*_*i*_ = 1, and homozygous derived in individual *i* if *v*_*i*_ = 2. So the DGS configuration produced by a mutation on such an edge is

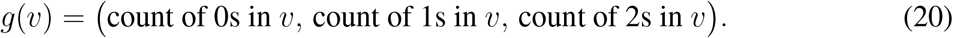

The calculation can then be treated as a recursion for which we must just do the bookkeeping. This has three steps: assign each coalescent edge a descendant vector, add waiting-time-weighted edge length to the DGS bin associated with that vector, and average over all possible next mergers up to the MRCA. The following formal recursion is this bookkeeping written out.

Let *S* = (*v*^(1)^, …, *v*^(*m*)^) be the current indexed collection of descendant vectors, with one entry for each slow lineage escaping from the fast phase. Thus *m* = |*S*| is the number of slow lineages. For convenience, time is measured in standard coalescent units, so that each pair of slow lineages coalesces at rate 1. Since there are 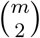 possible pairs of slow lineages, the total rate of coalescence is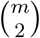, and the expected waiting time until the next coalescence is

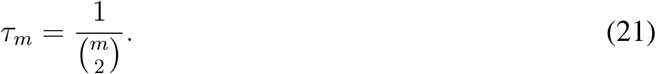

During this waiting time, each lineage contributes edge length to the specific DGS bin determined by that lineage’s descendant vector. Then one pair of lineages is chosen uniformly and merged.

Let 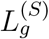 be the expected total edge length that would produce DGS configuration *g*, starting from state *S*. This value can be computed recursively by separating what happens before the next coalescence from what happens after it.

Before the next coalescence, the process remains in state *S* for an expected time *τ*_*m*_. During this waiting time, each lineage with descendant vector *v*^(*α*)^ continues to contribute edge length. As before, it contributes to 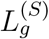 only if a mutation on that lineage would produce DGS configuration *g*. The total expected edge length contribution before the next coalescence is therefore

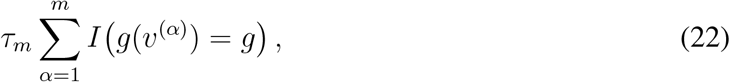

which adds up the number of extant edges that would contribute to each DGS bin, and scales them all by *τ*_*m*_.

After the waiting time, one pair of lineages coalesces. Since each pair is equally likely to be the next pair to coalesce, each of the 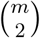 pairs has probability 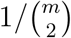. If the lineage instances indexed by *α* and *β* coalesce, the new state is *S*_*α,β*_, which we get by replacing *v*^(*α*)^ and *v*^(*β*)^ with their summed descendant vector. The expected future contribution after this coalescence is then 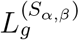. Averaging over all of the possible pairs of lineage instances gives

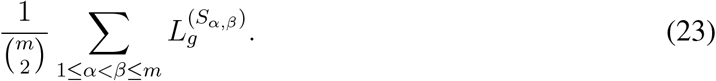

Adding the immediate contribution in Equation 22 to the expected future contribution in Equa-tion 23 gives

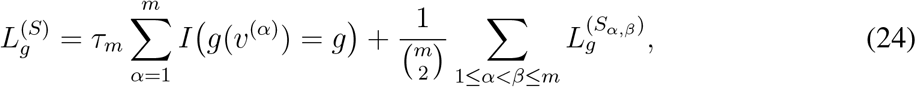

which is simply the sum of the two expressions. The recursion stops when *m* = 1:

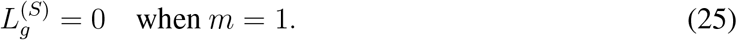

At that point all sampled homologs have coalesced into a single ancestral lineage, and a mutation above the sample MRCA would be carried by every sampled homolog and be monomorphic in the sample.

The last step is to marginalize over the fast-stage selfing outcomes defined above. Each fast-stage outcome **c** determines an initial slow state *S*(**c**). To briefly illustrate this, if *c*_*i*_ = 1, individual *i* contributes one slow lineage whose descendant vector has *v*_*i*_ = 2. If *c*_*i*_ = 0, individual *i* contributes two slow lineages, each with *v*_*i*_ = 1. In both cases, all other entries of those descendant vectors are zero.

The expected edge length producing DGS configuration *g* is finally computed by:

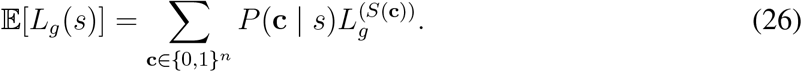

This expression sums over all 2^*n*^ binary fast-stage outcome vectors, which determine the starting states for the coalescent, and for each fast-stage outcome walks through the slow Kingman recursion for 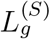.

### 5.1 Combinatorial approach

The recursion above gives an intuitive construction of the expected DGS edge lengths, but it is not very efficient because it enumerates every possible coalescent history. A key to making computation faster is recognizing that for the purposes of the DGS summation, the lineage labels are exchangeable. Specifically, fast-stage outcomes that differ only in the identities of the collapsed diploids (but are identical in the number of collapses) have the same DGS edge-length vector.

Let *x* be the number of collapsed diploids. The probability of this specific fast-stage group of outcomes is

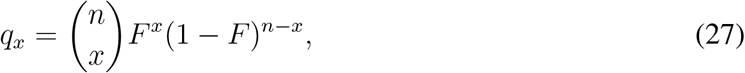

and the slow coalescent starts with

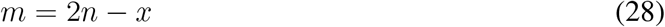

lineages.

A key observation is that for the *m*-lineage Kingman coalescent, the expected total length of all edges subtending exactly *j* of the initial lineages is

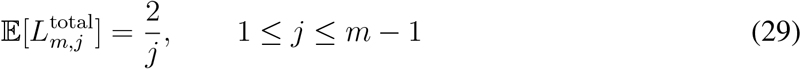

(Fu, 1995; Griffiths and Tavaré, 1998; Wakeley, 2009). Because the 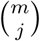 specified subsets of size *j* all behave the same under the coalescent, the expected length ancestral to any one specified subset is

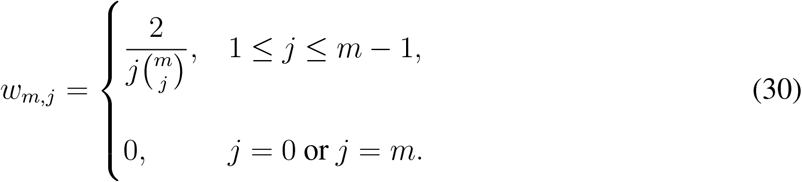

We can ignore the cases *j* = 0 and *j* = *m*, which correspond to empty and full descendant subsets, respectively, and do not produce polymorphic configurations.

We can now sort these expected edge lengths into their DGS contributions. Within a fast-stage outcome class containing *x* collapsed and *u* = *n* − *x* uncollapsed diploids, consider the subset of initial slow lineages subtended by an edge. Let *y* be the number of the *x* collapsed lineages included in the subset, *h* be the number of uncollapsed diploids represented by one homolog, and *d* be the number of uncollapsed diploids represented by both homologs. Such a subset contains

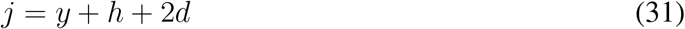

slow lineages. For a given number of collapsed individuals in the fast stage, the number of specified subsets with the same (*y, h, d*) description is

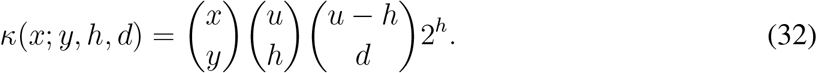

A mutation falling on an edge subtending this subset would contribute to the DGS configuration

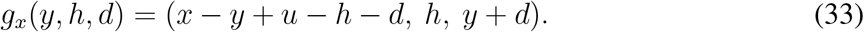

So, the edge-length vector for a particular fast-stage class can be written by setting *u* = *n* – *x* and *m* = 2*n* − *x*, and then looping over *y* = 0, …, *x, h* = 0, …, *u*, and *d* = 0, …, *u* − *h*. For each combination, we calculate *j* = *y* + *h* + 2*d*, skip the cases *j* = 0 and *j* = *m*, and add

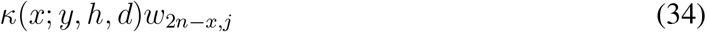

to the edge-length entry for its corresponding DGS configuration. To write this explicitly, the expected edge length yielding *g*, given the fast-stage class of exactly *x* collapsed diploids, is then

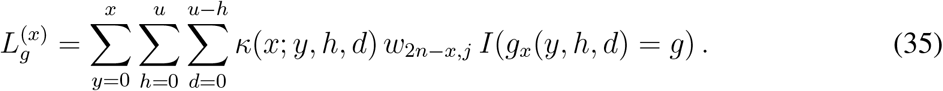

Finally, we repeat this calculation for *x* = 0, …, *n* and weight each class edge-length vector by its fast-stage probability *q*_*x*_. Our final expectation is then

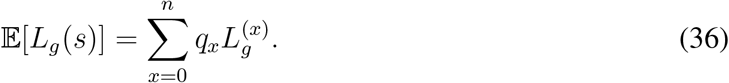

## 6 A three-diploid example

For *n* = 3 diploid individuals, there are eight polarized polymorphic DGS configurations:

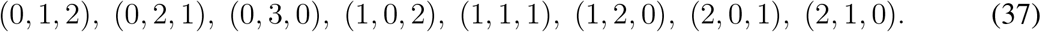

At selfing rate *s* = 0.7, the probability of coalescence during the fast phase is

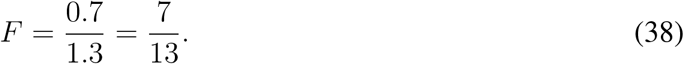

Evaluating the expected edge lengths (using the full recursion or through the combinatorial calcu-lation) gives the following probabilities:

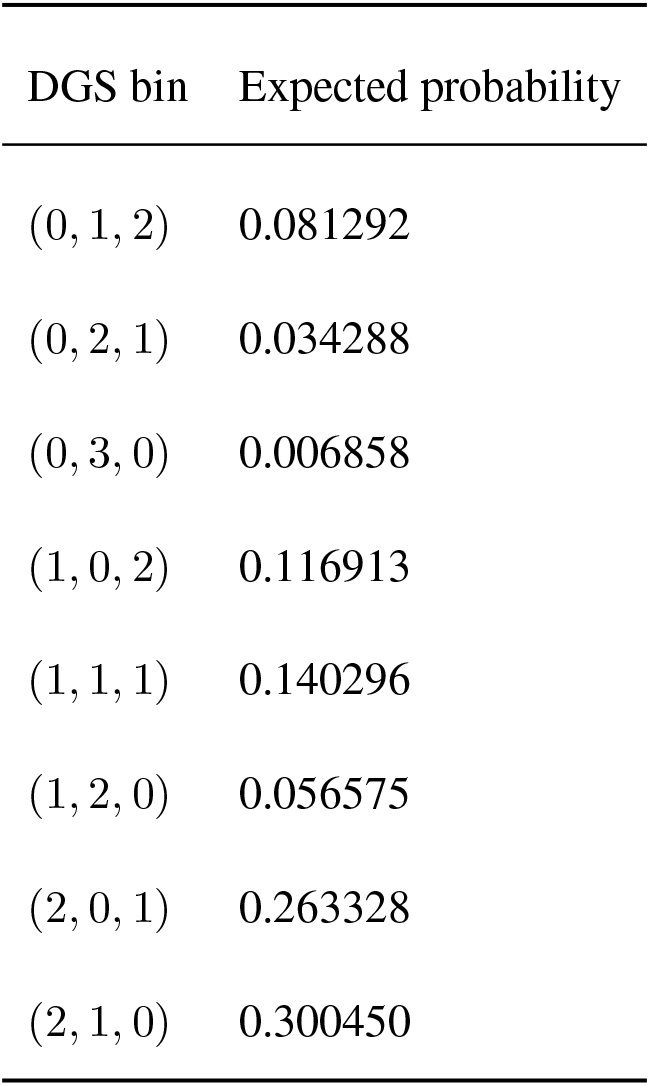

A full walk-through of the computation for 3 diploids is given in the supplementary notes: Supplementary Note 1 defines all variables used, Supplementary Note 2 follows the explicit recursion, and Supplementary Note 3 gives the faster combinatorial calculation.

## 7 Software implementation

We implemented these calculations in `selfdgs`, a Python package for constructing and fitting the DGS. `selfdgs` reads diploid, biallelic records from VCF files and can construct the DGS using various sources of ancestral-state information or by forming a folded DGS. For folded analyses, configurations related by reversal of ancestral vs. derived states are combined, and their model probabilities are summed. DGS configuration probabilities are evaluated by the combinatorial calculation described above, and the explicit state-by-state recursion is also an option as an independent reference.

`selfdgs` estimates the selfing rate by evaluating the conditional composite likelihood over a grid of possible selfing rates, *s*, and writes the DGS counts, likelihood curve, fit summary, and various diagnostics. It also supports reproducible fixed-size subset draws from user-defined sample groups and includes an optional workflow for validation with SLiM.

## 8 Simulation check

We used forward simulations in SLiM v5.2 (Haller et al., 2026) to test whether the analytical DGS probabilities match genotype configurations simulated under a Wright-Fisher model with partial selfing. We simulated true selfing rates in 5% increments

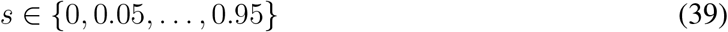

for census diploid population sizes

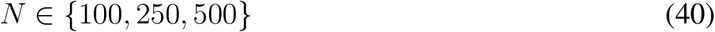

and samples of *n* = 3 or *n* = 6 diploid individuals, for a total of 20 * 3 * 2 = 120 parameter combinations. For every combination of *N* and *s*, we ran 10,000 independent neutral simulations in SLiM, each simulating a 1-kb locus, for a total of 10 Mb of simulated sequence per combination. The mutation rate was *µ* = 10^−7^ per bp per generation and the recombination rate was *r* = 5×10^−8^ per bp per generation. Each population was initialized with *N* diploids, maintained at constant size and selfing rate through cycle 10*N*, and sampled during the late stage of the last cycle. We drew *n* individuals without replacement from each simulated locus and recorded the DGS configuration of each segregating site. The *n* = 3 and *n* = 6 analyses used the same simulations, but the draws of individuals from each simulation happened independently. Because ancestral and derived states are known in the simulations, every segregating site was polarized.

For each parameter combination, DGS counts were pooled across the 10,000 loci before fitting, producing one selfing-rate estimate per parameter combination. Each locus came from a separate population with newly sampled individuals. Because of this, the simulations do not test how estimates vary among samples when the same individuals are used at every locus (which is useful for validating the theory, but for practical inference we think that shared individual selfing histories and the resulting identity disequilibrium may be an important source of sampling variance, particularly for small samples). We then evaluated the conditional composite log-likelihood

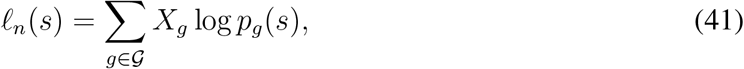

across candidate selfing rates on a grid of *s* = 0, 0.05, …, 0.95 individually for each parameter combination.

We found that we can efficiently calculate the DGS probabilities for any defined selfing rate (shown for *s* = 0.7 with sampling *n* = 3 in Figure 2A and *n* = 6 in Figure 2D). These analytical probabilities closely match the observed DGS frequencies calculated from the SLiM simulations across a range of population sizes (Figure 2B for *n* = 3 and Figure 2E for *n* = 6). Further, we calculated the relative conditional composite log-likelihood, *l*_*n*_(*s*) − max *l*_*n*_(*s*), across the candidate selfing-rate possibilities and found that the likelihood is maximized near the true simulated value of *s* = 0.7 in all cases (Figure 2C and F). These results show that the analytical probabilities match the forward simulations and that the calculation extends from the three-diploid worked example to six diploids.

**Fig 2.**
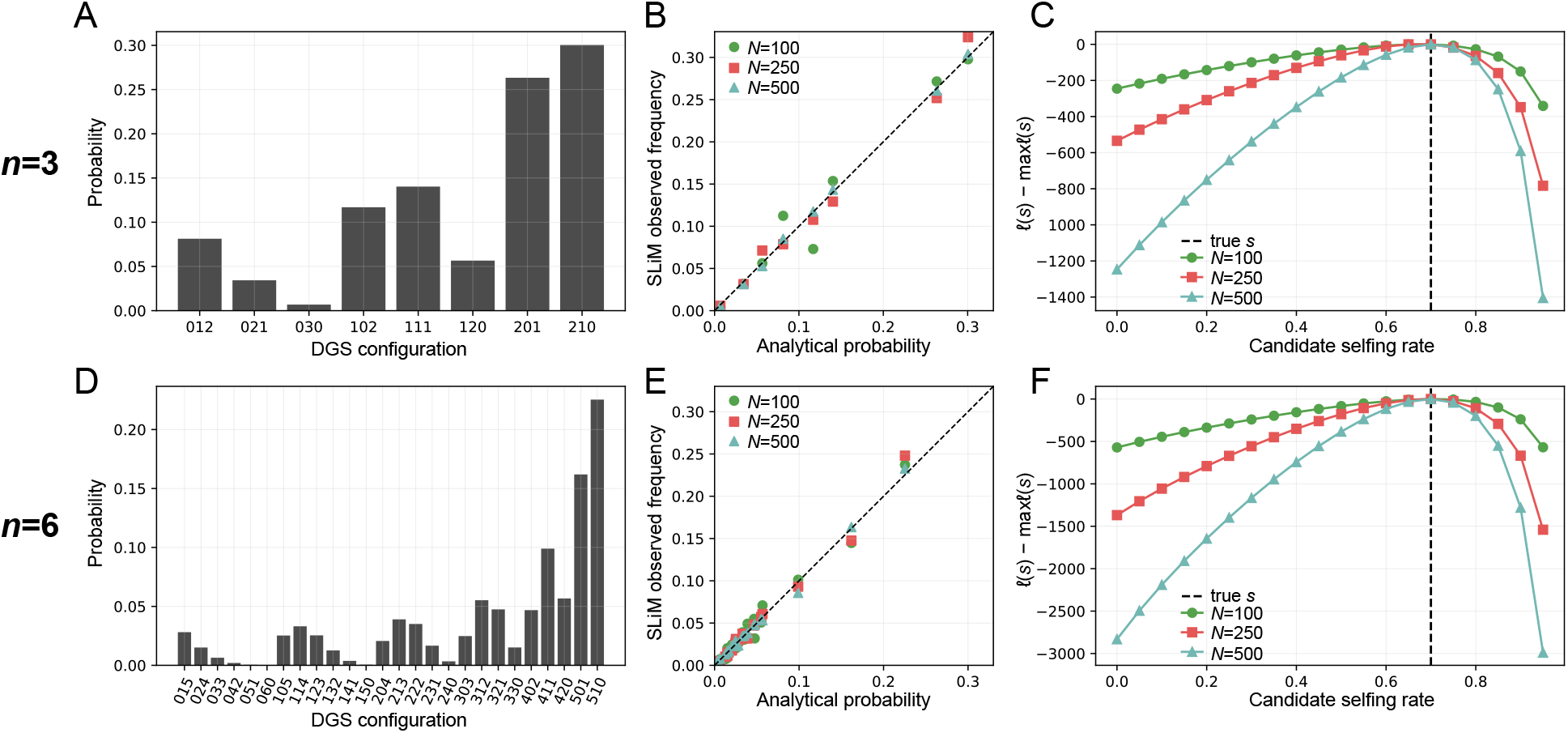
Analytical and simulated DGS probabilities for samples of three and six diploid individuals under partial selfing at *s* = 0.7. The top row shows results for *n* = 3 sampled diploids and the bottom row shows results for *n* = 6 sampled diploids. (A,D) Expected DGS probabilities for *s* = 0.7. Axis labels concatenate the counts (*n*_0_, *n*_1_, *n*_2_); for example, 012 denotes (0, 1, 2). (B,E) Observed DGS frequencies from SLiM simulations plotted against analytical probabilities. Each point is one DGS configuration, and colors/shapes indicate census population size (*N* = 100, 250, 500). For *n* = 3, the simulations contain 534, 1, 207, and 2, 444 segregating sites for *N* = 100, 250, and 500, respectively. For *n* = 6, the simulations contain 691, 1, 762, and 3, 456 segregating sites for *N* = 100, 250, and 500, respectively. The dashed line shows equality. (C,F) Relative composite log-likelihood, *l*_*n*_(*s*) − max *l*_*n*_(*s*), evaluated along a grid of candidate selfing rates using DGS counts pooled across 10,000 independently simulated loci, with newly sampled individuals at each locus. The dashed vertical line marks the true simulated value, *s* = 0.7.

We next used the full simulation grid to examine the composite-likelihood surface across true and candidate selfing rates (Figure 3). For each simulated rate, we calculated the likelihood profile relative to its maximum over the candidate-*s* grid. For visualization, relative log-likelihood values below −80 were clipped.

**Fig 3.**
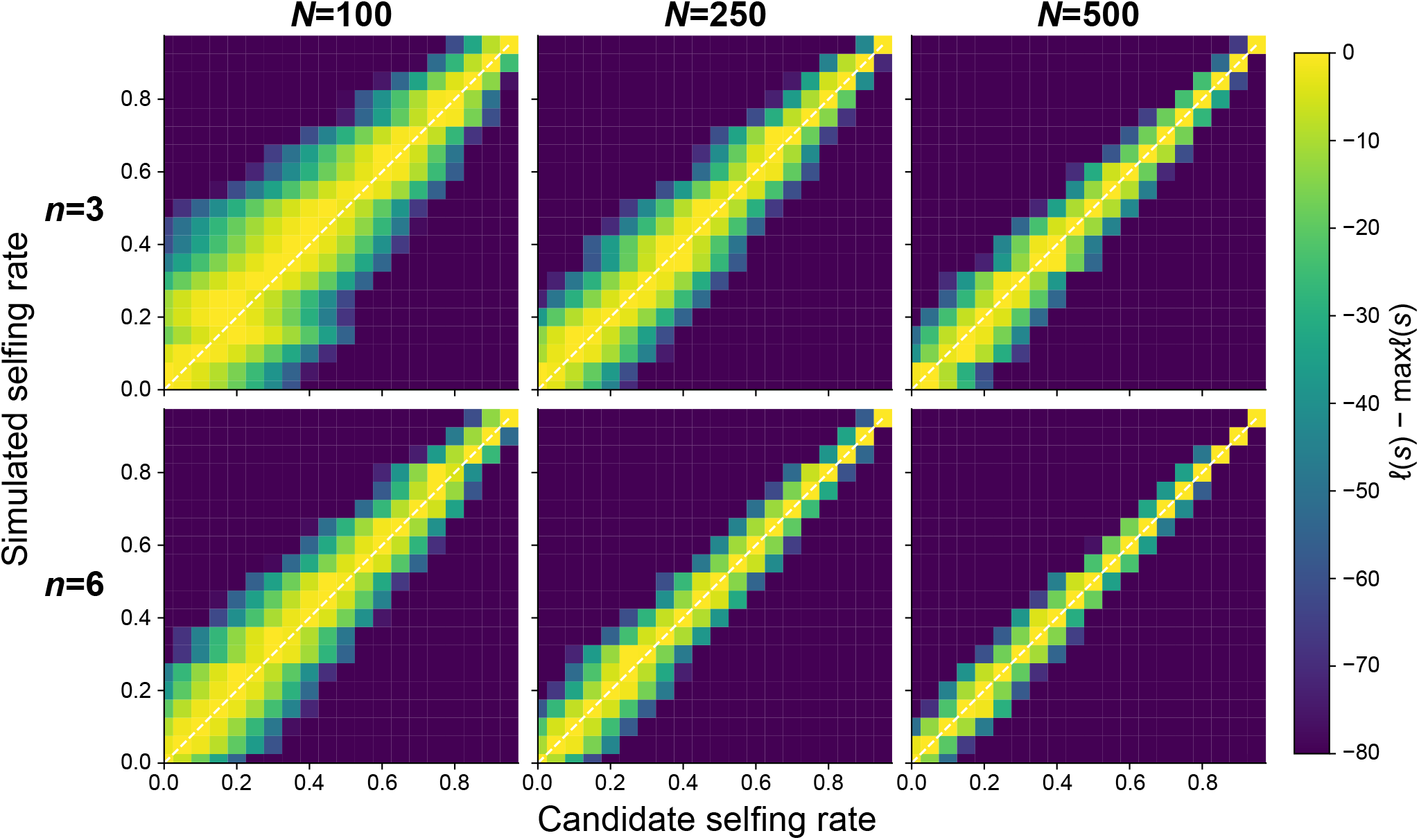
Composite-likelihood surfaces comparing simulated vs. candidate selfing rates. Top and bottom panel rows correspond to samples of *n* = 3 and *n* = 6 diploid individuals, respectively, and columns show census population sizes *N* = 100, 250, and 500. The x-axis indicates the candidate selfing rate *s*, and the vertical axis denotes the true selfing rate used in the forward simulation. For each simulated rate, colors show the relative conditional composite log-likelihood, *l*_*n*_(*s*) − max *l*_*n*_(*s*), evaluated over the candidate-rate grid. Yellow indicates the best-scoring rates, and darker colors indicate lower support. Values below −80 are displayed at the lower limit of the color scale. The dashed white line is the one-to-one relationship between the simulated and candidate rates. Concentration of the likelihood near this line indicates that the DGS likelihood recovers the simulated selfing rate under this sampling design. The bright region shows the range of candidate selfing rates with relatively high composite likelihoosd for each pooled dataset.

Across both sample sizes and the three population sizes, the region of greatest support follows the one-to-one relationship between the simulated and candidate selfing rates, marked by the dashed diagonal in Figure 3. The likelihood surface is broadest for *N* = 100 and becomes more concentrated around the diagonal as population size (and therefore the number of segregating sites) increases. It also becomes more concentrated at higher selfing rates. The same overall pattern occurs for samples of three and six diploids.

## 9 Discussion

Reliable estimates of population self-fertilization rates are important for interpreting the evolutionary consequences of selfing and for comparing mating systems across taxa, yet such estimates remain unavailable for many species (Goodwillie et al., 2005; Meyer et al., 2026). Selfing presents an evolutionary trade-off. Previous work suggests it should be favored by selection under mate limitation, in which case it helps ensure reproductive success. However, selfing also skews genotype frequencies, depletes genome-wide diversity, and potentially reduces the adaptive capacity and long-term persistence of populations (Busch and Delph, 2012; Cheptou, 2019; Glémin and Ronfort, 2013; Shimizu and Tsuchimatsu, 2015; Wright et al., 2013). Here we develop the DGS as a compact genomic summary for estimating population selfing rates from diploid SNP data, based on the standard coalescent model of partial selfing and implemented in `selfdgs` (Nordborg and Donnelly, 1997).

Selfing changes how alleles are paired within individuals, altering the relative frequencies of diploid genotype configurations and thus leaving a genomic signature (David et al., 2007). The DGS captures this information while remaining simple to construct from diploid SNP data, e.g. genotype calls stored in standard VCF files. We derived its expected distribution under an arbitrary selfing rate using two equivalent approaches: an intuitive recursive calculation and a more computationally efficient combinatorial calculation, the latter of which is used by default in `selfdgs`.

The simulations examined here demonstrate close agreement between analytical expectations and observed DGS frequencies from simulations.

By summarizing per-site genotype configurations while ignoring linkage, phase, and genealogy information, the DGS retains the full distribution of diploid genotype configurations across sites rather than reducing these data to a single inbreeding statistic like *F*_*IS*_, while avoiding the complexity of methods that model the inbreeding history of each sampled individual or the local genealogy of each locus (David et al., 2007; Redelings et al., 2015; Strütt et al., 2023). More specifically, because the DGS probabilities depend on the selfing rate only through *F* = *s/*(2 − *s*), DGS-based estimation targets the same equilibrium inbreeding parameter that underlies *F*-based estimates of selfing. The distinction is statistical more than biological. Estimators like Weir–Cockerham *F*_*IS*_ reduce genotype counts across loci to heterozygosity-based moments, while the DGS instead fits the full distribution of genotype configurations expected under the partial-selfing coalescent. So the full DGS holds on to configuration-specific information that is lost in a single *F*_*IS*_ estimate, including both how alleles are arranged among diploid genotypes at a given allele count and variation in the relative frequencies of allele-count classes.

DGS-based estimates are calculated under a neutral panmictic model, and so population structure, genotyping error, or other processes that might alter the pairing of alleles within individuals could shift DGS frequencies in ways that resemble selfing (Bürkli et al., 2017; Gao et al., 2007). Estimates should therefore be interpreted as the selfing rate that best reproduces the observed DGS under the fitted model rather than necessarily as the true mating-system selfing rate. Similarly, changes in population size or selfing rate through time could affect estimates, and we have not yet evaluated the robustness of inference with the DGS to these more complex models. Finally, the DGS is built from within-individual genotype calls and therefore shares the sensitivity of *F*_*IS*_ to allelic dropout and similar artifacts that score heterozygotes as homozygotes, as described in David et al. (2007). Complementary methods can address different confounders, with RMES (David et al., 2007) being less sensitive to genotyping artifacts that occur independently across loci, in-Struct (Gao et al., 2007) jointly modeling population structure, and teSMC (Strütt et al., 2023) leveraging information from recombination.

Additionally, linkage among sites could affect inference. Marginal DGS probabilities can be combined as a composite likelihood, but linked sites contribute less independent information than the DGS likelihood assumes, especially since selfing reduces the efficacy of recombination (Golding and Strobeck, 1980; Kogan et al., 2025; Padhukasahasram et al., 2008; Strütt et al., 2023). In empirical applications, LD pruning could reduce the effects of physical linkage. However, even unlinked loci share an individual’s recent history of selfing (Redelings et al., 2015). Sampling more loci from the same individuals does not sample new individual histories, so it may improve precision less than the likelihood curves suggest. Our simulations use newly sampled individuals at each locus and do not measure this effect. Testing precision for genomic samples will require samples from independent simulated populations, keeping each sample of individuals the same across loci. Uncertainty could in principle be quantified using the Godambe Information Matrix approach of Coffman et al. (2016), but this would need to account for dependence among loci and be tested with such samples.

The framework here is also extendable beyond diploids. For higher ploidy, descendant vectors could track allelic dosage within each sampled individual, although the within-individual ancestral process and associated combinatorics would be more complex. This possibility supports a broader class of genotype-configuration spectra that retain information about how alleles are arranged within individuals (Ferretti et al., 2018).

Here we establish the expected DGS under the standard partial-selfing coalescent, providing a theoretical foundation for estimating selfing rates from population genomic data. By linking diploid genotype variation to an explicit genealogical model, the DGS offers a practical and theoretically grounded way of studying the distribution and evolution of selfing across species.

## 10 Data Availability

The software package `selfdgs` (https://github.com/pmckenz1/selfdgs) contains code for constructing and performing inference under the DGS. The simulation analyses for making the figures here are preserved in the analysis repository (https://github.com/pmckenz1/dgs_manuscript).

## 11 Acknowledgements

We are grateful to John Wakeley for feedback on an early draft. This project was inspired by discussions at the 2025 Speciation Gordon Research Conference.

## Supplementary Note 1: Reference guide to notation

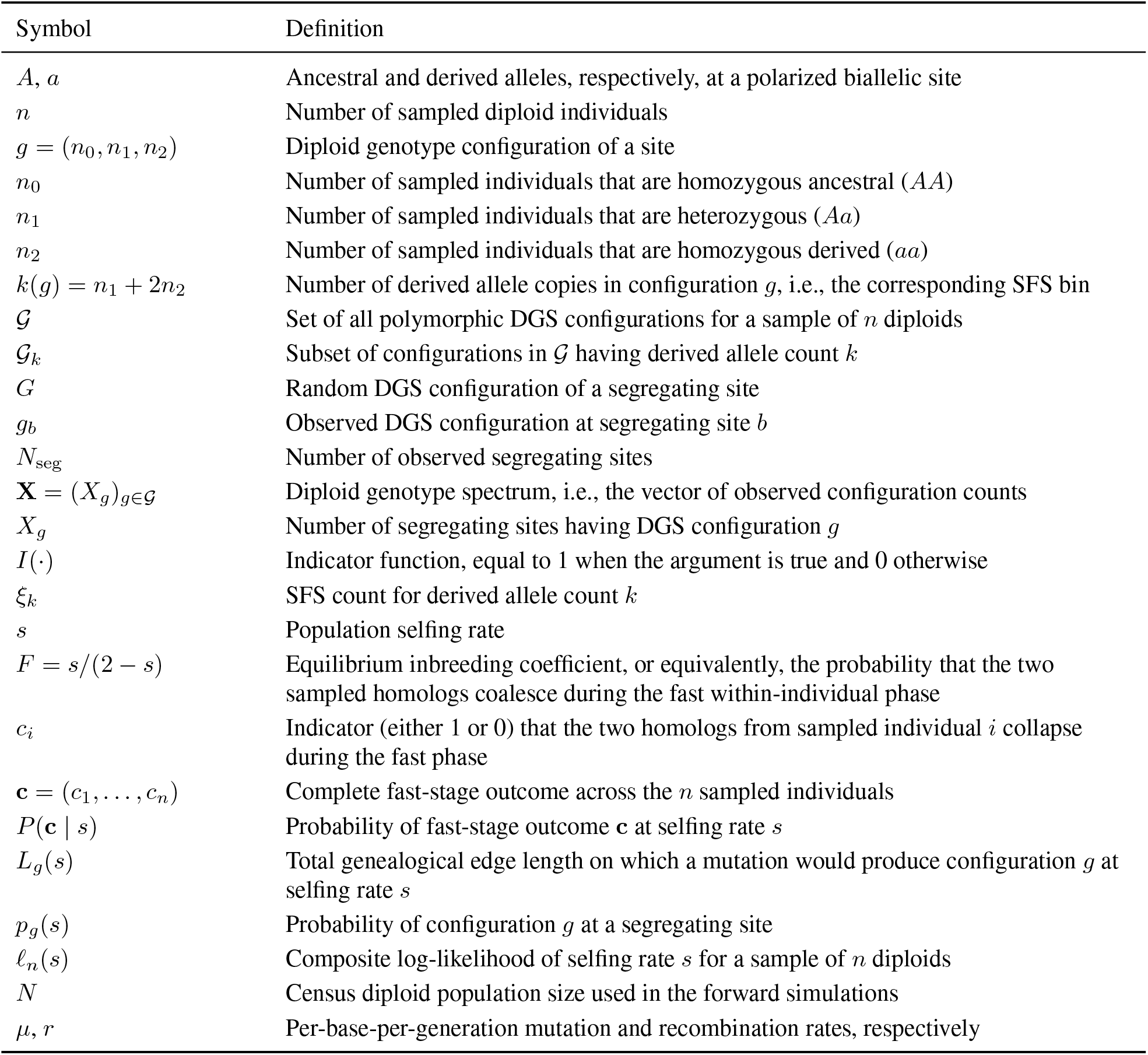

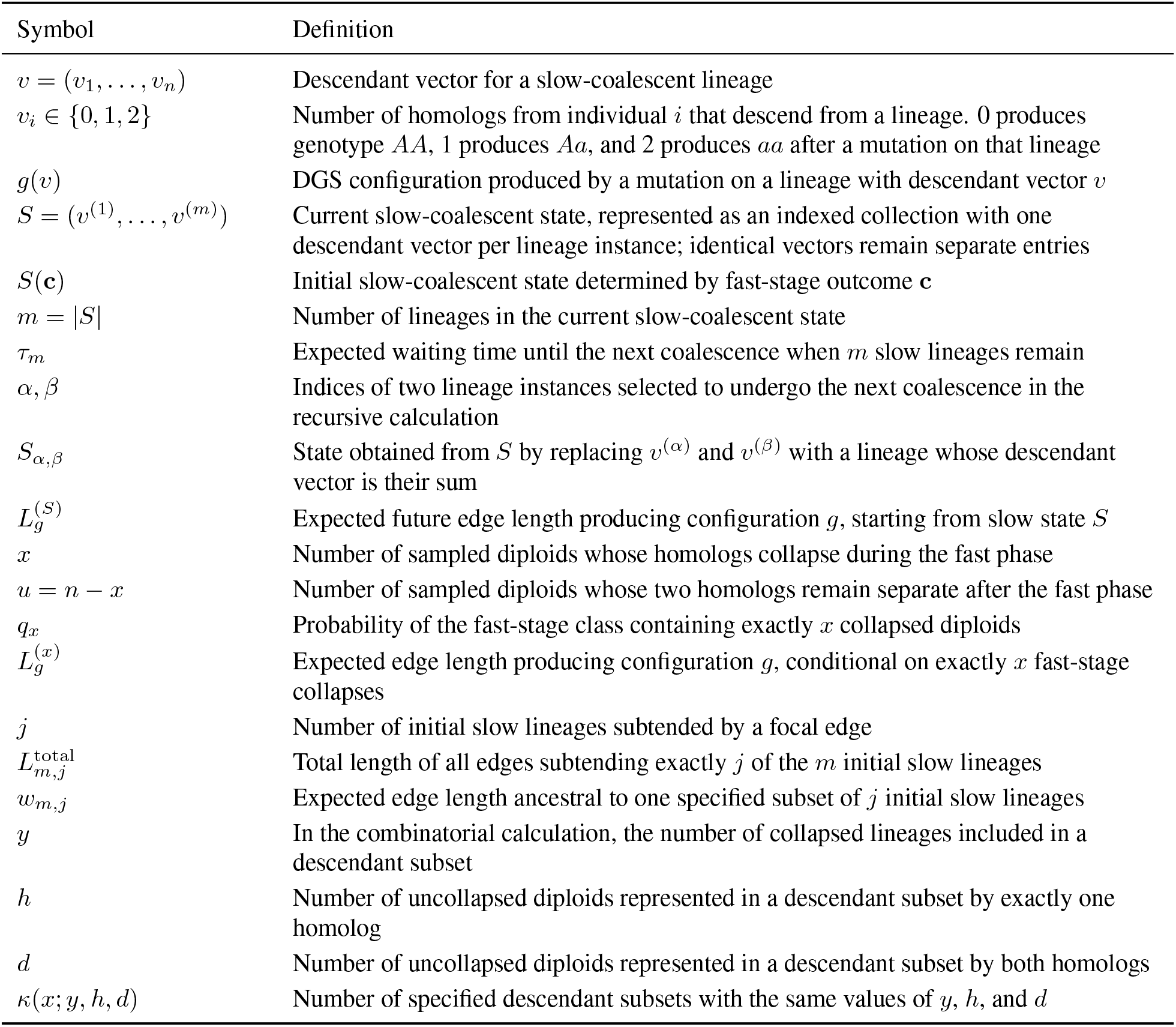

## Supplementary Note 2: Explicit coalescent-state derivation for s=0.7 example

### S1. Purpose and notation

This is a fully worked example showing how to compute the expected relative frequencies of diploid genotype spectrum (DGS) bins for a sample of three diploid individuals under the standard fast-selfing approximation of Nordborg and Donnelly (1997) using the recursion method. The goal is to marginalize over the possible fast-stage outcomes and their possible slow Kingman coalescent histories.

For a segregating mutation, a DGS bin is written as

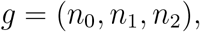

where *n*_0_ is the number of homozygous ancestral individuals, *n*_1_ is the number of heterozygous individuals, and *n*_2_ is the number of homozygous derived individuals. In the tables below, we will write the bin (*n*_0_, *n*_1_, *n*_2_) more compactly as *n*_0_*n*_1_*n*_2_, just for convenience. For example, 201 means (2, 0, 1).

We sample three diploid individuals. The six homologs are labeled 0 through 5:

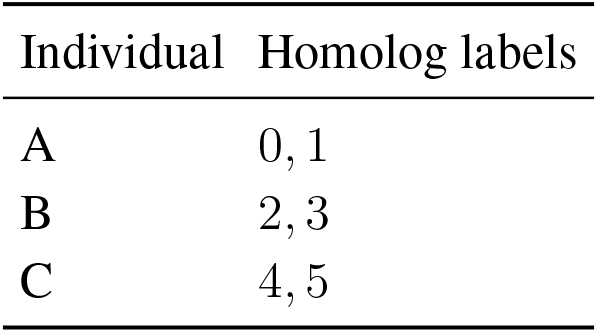

| Individual | Homolog labels |
| --- | --- |
| A | 0, 1 |
| B | 2, 3 |
| C | 4, 5 |

An edge is assigned to a DGS bin by asking how many sampled homologs from each diploid individual descend from that edge. So for example, a mutation on edge (01) is present in both homologs of individual A and absent from individuals B and C, so it generates the DGS bin (2, 0, 1), written 201. A mutation on edge 4 is present in one homolog of individual C only, so it generates (2, 1, 0), written 210.

All edge lengths below are in coalescent units (the usual time scale of (2 − *s*)*N* generations is common to all of the slow lineages and cancels when we divide by the total edge length to get relative DGS probabilities). We stop the calculation at the sample MRCA because mutations occurring earlier would be present in every homolog (i.e. the site would be monomorphic).

### S2. Fast-stage outcomes

Under partial selfing, the two homologs sampled from the same diploid go through a fast within-individual phase. With selfing rate *s* = 0.7, the inbreeding coefficient is

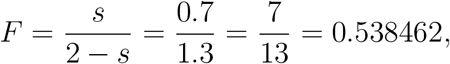

and

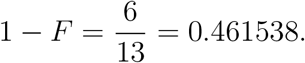

Each sampled diploid pair collapses in the fast phase with probability *F* or might remain as two distinct slow lineages with probability 1 − *F*. So a particular outcome with *x* collapsed diploids for this example has probability

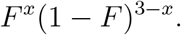

| Fast outcome class | Slow-lineage starting state | Probability | Decimal |
| --- | --- | --- | --- |
| 0 collapsed | 0 1 2 3 4 5 | $(1 - F)^3 = 216/2197$ | 0.098316 |
| 1 collapsed | (01) 2 3 4 5 | $F(1 - F)^2 = 252/2197$ | 0.114702 |
| 1 collapsed | 0 1 (23) 4 5 | $F(1 - F)^2 = 252/2197$ | 0.114702 |
| 1 collapsed | 0 1 2 3 (45) | $F(1 - F)^2 = 252/2197$ | 0.114702 |
| 2 collapsed | (01) (23) 4 5 | $F^2(1 - F) = 294/2197$ | 0.133819 |
| 2 collapsed | 0 1 (23) (45) | $F^2(1 - F) = 294/2197$ | 0.133819 |
| 2 collapsed | (01) 2 3 (45) | $F^2(1 - F) = 294/2197$ | 0.133819 |
| 3 collapsed | (01) (23) (45) | $F^3 = 343/2197$ | 0.156122 |

### S3. The recursion

The slow phase is a Kingman coalescent on the lineages that remain after the fast phase. If the current slow state has *m* lineages, the expected waiting time until the next coalescence is

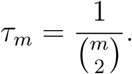

During this waiting time, every current lineage contributes *τ*_*m*_ units of edge length to its associated DGS bin.

Afterward, one pair of lineages is chosen uniformly to merge. So, if *m* = 3, there are three possible next mergers, and each has probability 1*/*3 (these are the Sub1, Sub2, and Sub3 rows below). Their edge-length contributions are multiplied by 1*/*3 because they are conditional alternatives (only one of the three substates is reached). Similarly, when *m* = 4, there are six possible first mergers (4 choose 2), so the six option totals are averaged with weight 1*/*6.

Represent the slow state as an indexed collection *S* = (*v*^(1)^, …, *v*^(*m*)^), with one entry for each lineage instance. This keeps distinct lineages separate even when their descendant vectors are identical. Let 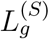 be the expected future edge length in DGS bin *g*, starting from slow state *S*. If |*S*| = *m*, then

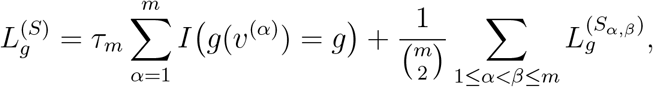

where 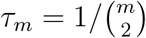, *S*_*α,β*_ is the state formed by merging the lineage instances indexed by *α* and *β*, and 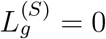 once only one lineage remains.

### S4. Worked recursion for one fast outcome: (01) (23) 4 5

As an example, we can look at the full recursion for the fast-stage outcome in which individuals A and B collapse, but individual C does not. The starting slow state is

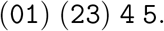

There are initially four slow lineages. The expected waiting time before the next coalescence is

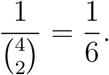

The current edges contribute as follows:

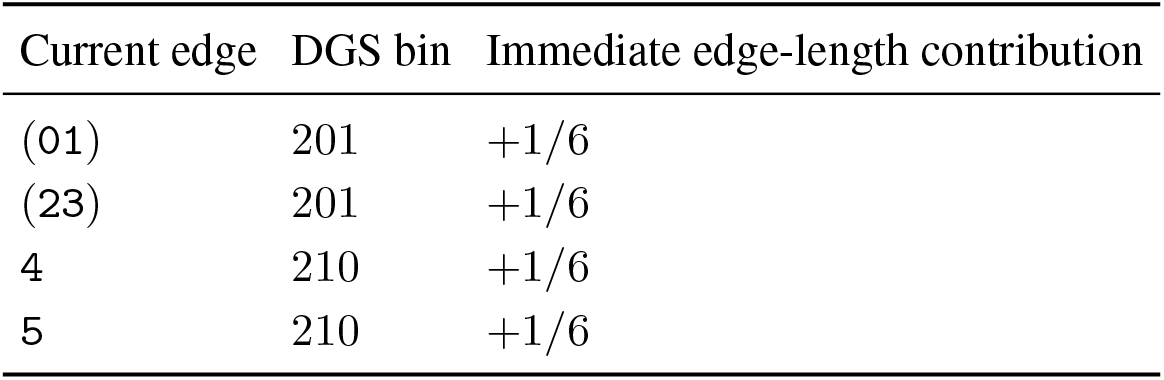

These initial 1*/*6 contributions are held aside and added only after averaging over the six possible next mergers. The six possible pairwise mergers from the four-lineage state are the six options below. In each option table, the “Immediate” rows are the edge lengths accumulated in the three-lineage state itself, while the “Sub1/Sub2/Sub3” rows are the three possible two-lineage states produced by the next coalescence. Because these substates are alternative outcomes, their contributions are averaged with weight 1*/*3 in the total for each option (lower table).

**Option 1:** (0123) 4 5.

| Recursion step | Edge | DGS bin | Contribution |
| --- | --- | --- | --- |
| Immediate state; time = $1/\binom{3}{2} = 1/3$ | (0123) | 102 | $+1/3$ |
| Immediate state; time = $1/\binom{3}{2} = 1/3$ | 4 | 210 | $+1/3$ |
| Immediate state; time = $1/\binom{3}{2} = 1/3$ | 5 | 210 | $+1/3$ |
| Sub1: (01234) 5; time = $1/\binom{2}{2} = 1$ | (01234) | 012 | $+1$ |
| Sub1: (01234) 5; time = $1/\binom{2}{2} = 1$ | 5 | 210 | $+1$ |
| Sub2: (01235) 4; time = $1/\binom{2}{2} = 1$ | (01235) | 012 | $+1$ |
| Sub2: (01235) 4; time = $1/\binom{2}{2} = 1$ | 4 | 210 | $+1$ |
| Sub3: (0123) (45); time = $1/\binom{2}{2} = 1$ | (0123) | 102 | $+1$ |
| Sub3: (0123) (45); time = $1/\binom{2}{2} = 1$ | (45) | 201 | $+1$ |

| DGS bin | Calculation | Total |
| --- | --- | --- |
| 012 | $\frac{1}{3}(1 + 1)$ | $2/3$ |
| 102 | $\frac{1}{3} + \frac{1}{3}(1)$ | $2/3$ |
| 201 | $\frac{1}{3}(1)$ | $1/3$ |
| 210 | $\frac{1}{3} + \frac{1}{3} + \frac{1}{3}(1 + 1)$ | $4/3$ |

**Options 2**. (014) (23) 5.

**Option 2:** (014) (23) 5.
| Recursion step | Edge | DGS bin | Contribution |
| --- | --- | --- | --- |
| Immediate state; time = $1/\binom{3}{2} = 1/3$ | (014) | 111 | $+1/3$ |
| Immediate state; time = $1/\binom{3}{2} = 1/3$ | (23) | 201 | $+1/3$ |
| Immediate state; time = $1/\binom{3}{2} = 1/3$ | 5 | 210 | $+1/3$ |
| Sub1: (01234) 5; time = $1/\binom{2}{2} = 1$ | (01234) | 012 | $+1$ |
| Sub1: (01234) 5; time = $1/\binom{2}{2} = 1$ | 5 | 210 | $+1$ |
| Sub2: (0145) (23); time = $1/\binom{2}{2} = 1$ | (0145) | 102 | $+1$ |
| Sub2: (0145) (23); time = $1/\binom{2}{2} = 1$ | (23) | 201 | $+1$ |
| Sub3: (014) (235); time = $1/\binom{2}{2} = 1$ | (014) | 111 | $+1$ |
| Sub3: (014) (235); time = $1/\binom{2}{2} = 1$ | (235) | 111 | $+1$ |

| DGS bin | Calculation | Total |
| --- | --- | --- |
| 012 | $\frac{1}{3}(1)$ | $1/3$ |
| 102 | $\frac{1}{3}(1)$ | $1/3$ |
| 111 | $\frac{1}{3} + \frac{1}{3}(2)$ | $1$ |
| 201 | $\frac{1}{3} + \frac{1}{3}(1)$ | $2/3$ |
| 210 | $\frac{1}{3} + \frac{1}{3}(1)$ | $2/3$ |

**Options 3-5**. Options 3-5 have the same option totals as Option 2 by relabeling sampled homologs (this exchangeability is leveraged in the combinatorial approach in Supplementary Note 3):

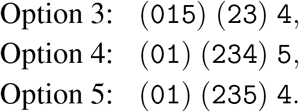

Each has option totals 012 = 1/3, 102 = 1/3, 111 = 1, 201 = 2/3, and 210 = 2/3.

**Option 6:** (01) (23) (4 5).

| Recursion step | Edge | DGS bin | Contribution |
| --- | --- | --- | --- |
| Immediate state; time = $1/\binom{3}{2} = 1/3$ | (01) | 201 | +1/3 |
| Immediate state; time = $1/\binom{3}{2} = 1/3$ | (23) | 201 | +1/3 |
| Immediate state; time = $1/\binom{3}{2} = 1/3$ | (45) | 201 | +1/3 |
| Sub1: (0123) (45); time = $1/\binom{2}{2} = 1$ | (0123) | 102 | +1 |
| Sub1: (0123) (45); time = $1/\binom{2}{2} = 1$ | (45) | 201 | +1 |
| Sub2: (0145) (23); time = $1/\binom{2}{2} = 1$ | (0145) | 102 | +1 |
| Sub2: (0145) (23); time = $1/\binom{2}{2} = 1$ | (23) | 201 | +1 |
| Sub3: (01) (2345); time = $1/\binom{2}{2} = 1$ | (01) | 201 | +1 |
| Sub3: (01) (2345); time = $1/\binom{2}{2} = 1$ | (2345) | 102 | +1 |

| DGS bin | Calculation | Total |
| --- | --- | --- |
| 012 | 0 | 0 |
| 102 | $\frac{1}{3}(1 + 1 + 1)$ | 1 |
| 111 | 0 | 0 |
| 201 | $\frac{1}{3} + \frac{1}{3} + \frac{1}{3} + \frac{1}{3}(1 + 1 + 1)$ | 2 |
| 210 | 0 | 0 |

### S5. Averaging over the six options for the fast-stage outcome (01) (23) 4 5

Because each pairwise merger from the four-lineage state is equally likely, the six option totals are averaged with weight 1*/*6. This gives the expected future edge length after the first four-lineage waiting interval, but before adding the initial 1*/*6 contributions from the four-lineage state itself.

| DGS bin | Average calculation | Average |
| --- | --- | --- |
| 012 | $(2/3 + 1/3 + 1/3 + 1/3 + 1/3 + 0)/6$ | $1/3$ |
| 102 | $(2/3 + 1/3 + 1/3 + 1/3 + 1/3 + 1)/6$ | $1/2$ |
| 111 | $(0 + 1 + 1 + 1 + 1 + 0)/6$ | $2/3$ |
| 201 | $(1/3 + 2/3 + 2/3 + 2/3 + 2/3 + 2)/6$ | $5/6$ |
| 210 | $(4/3 + 2/3 + 2/3 + 2/3 + 2/3 + 0)/6$ | $2/3$ |

Now add the initial four-lineage contributions. DGS bin 201 receives 1*/*6 + 1*/*6 = 1*/*3, and DGS bin 210 receives 1*/*6 + 1*/*6 = 1*/*3.

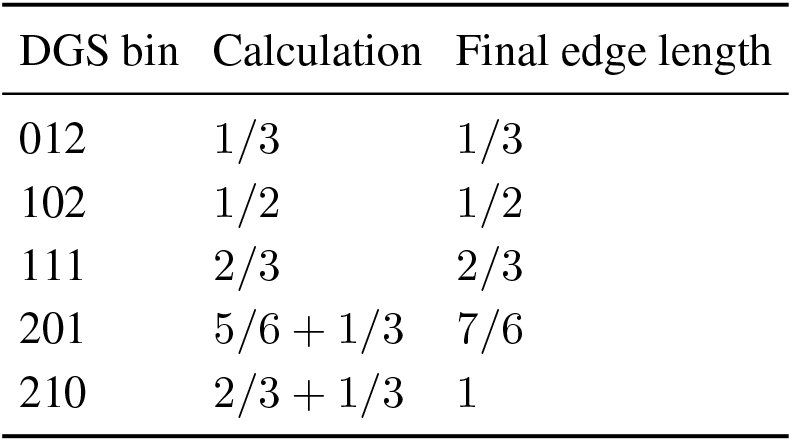

As a check,

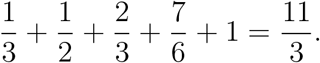

This equals the expected total edge length from a four-lineage Kingman state:

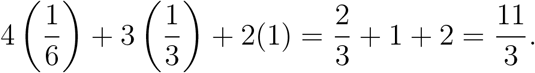

### S6. Edge lengths for all fast-stage outcomes

The same recursion is repeated for each fast-stage outcome. By symmetry, the three single-collapse outcomes have identical edge-length vectors, and the three double-collapse outcomes have identical edge-length vectors. Zeros indicate that no edge in that slow genealogy can generate the given DGS configuration.

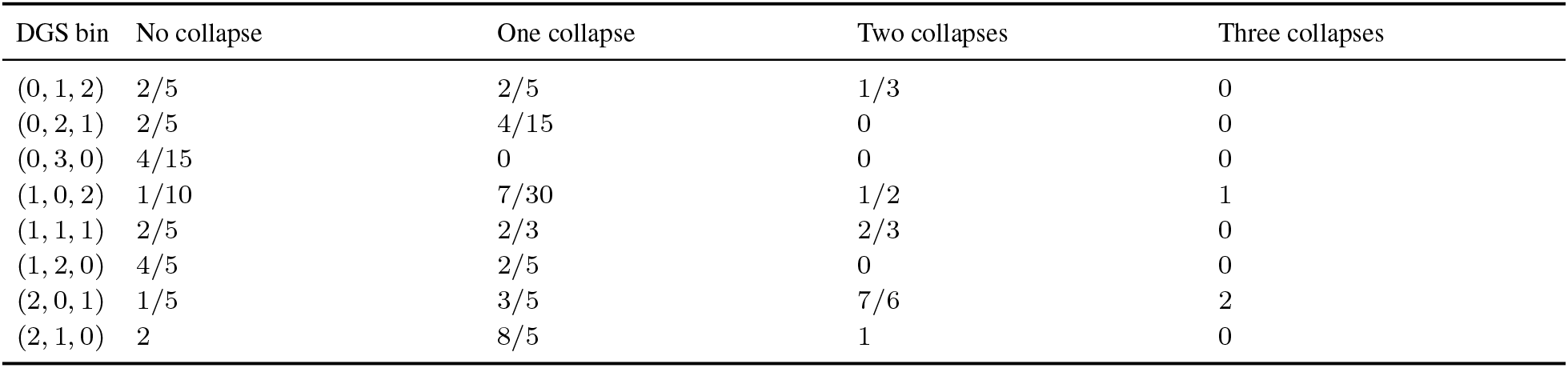

### S7. Weighted edge lengths across the entire fast-stage space

For each DGS bin *g*, we can weight the edge length by the probability of each fast-stage outcome (i.e., marginalizing over the possible fast stage outcomes). Let *p*_0_ be the probability of the no-collapse outcome, *p*_1_ the probability of any particular one-collapse outcome, *p*_2_ the probability of any particular two-collapse outcome, and *p*_3_ the probability of the three-collapse outcome. There are three one-collapse outcomes and three two-collapse outcomes, so the middle two terms are multiplied by 3:

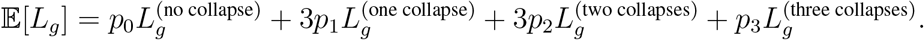

The fast-stage probabilities are

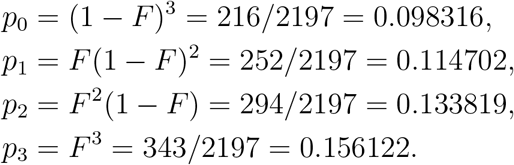

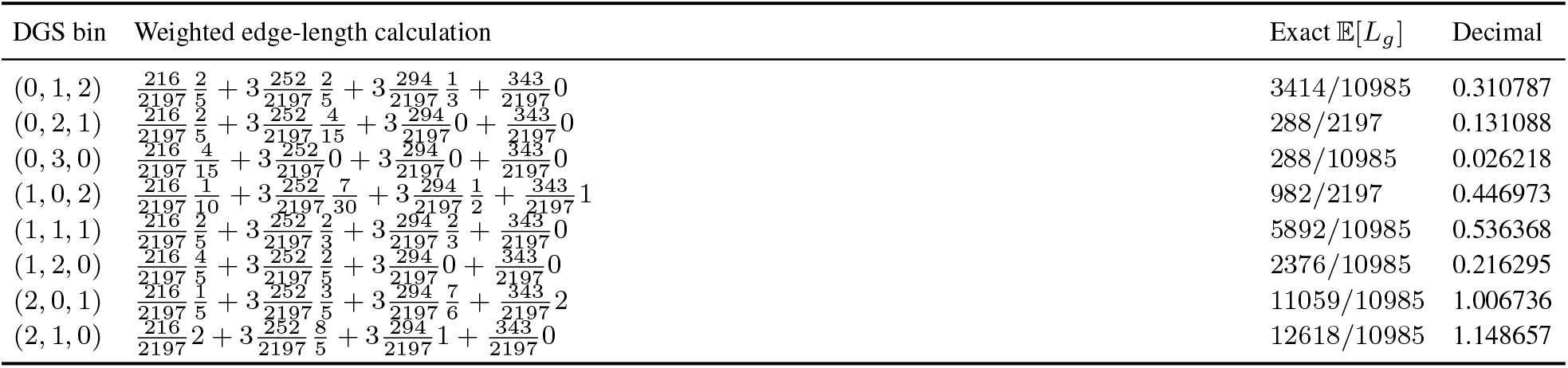

### S8. Final DGS probabilities

The total weighted edge length is

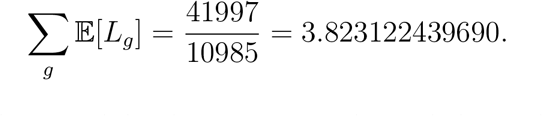

Dividing each weighted edge length by this total gives the probability that a segregating mutation falls in each DGS bin.

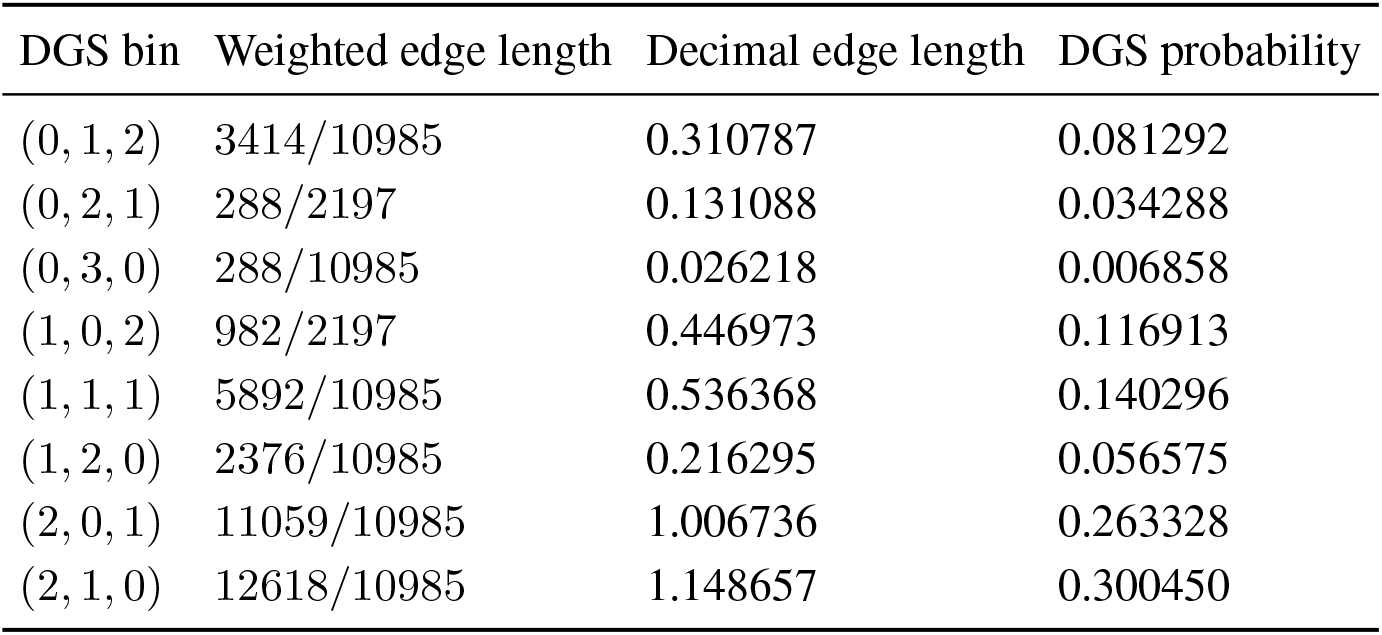

### S9. Interpretation and connection to the SFS

The largest DGS probabilities for *s* = 0.7 are for (2, 1, 0) and (2, 0, 1). These are configurations with a singleton derived allele or a single homozygous-derived individual, respectively. The enrichment of homozygous derived configurations is because of the high probability of fast selfing collapse. Once the two homologs from an individual collapse (backward in time), mutations on the resulting slow lineage are carried by both homologs in that individual.

The ordinary unfolded SFS would collapse DGS bins by derived allele count

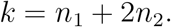

For this example, the projection to the SFS is:

| Derived allele count $k$ | DGS bins collapsed into this SFS bin | Probability |
| --- | --- | --- |
| 1 | (2, 1, 0) | 0.300450 |
| 2 | (1, 2, 0) + (2, 0, 1) | $0.056575 + 0.263328 = 0.319903$ |
| 3 | (0, 3, 0) + (1, 1, 1) | $0.006858 + 0.140296 = 0.147154$ |
| 4 | (0, 2, 1) + (1, 0, 2) | $0.034288 + 0.116913 = 0.151201$ |
| 5 | (0, 1, 2) | 0.081292 |

So this illustrates the advantage of using the DGS rather than only the SFS: two mutations with the same derived allele count can correspond to different diploid genotype arrangements. For example, the SFS doubleton bin *k* = 2 combines (1, 2, 0), where the two derived copies are in two heterozygotes, with (2, 0, 1), where the two derived copies are paired as one homozygous-derived individual. Those two configurations have very different expected probabilities under selfing.

## Supplementary Note 3: Efficient combinatorial derivation for s=0.7 example

### C1. Purpose and relationship to the full recursion

Here we will repeat the three-diploid calculation from Supplementary Note 2, but we will use the more efficient combinatorial approach implemented by default in `selfdgs`. The underlying biological model and the resulting DGS probabilities are unchanged, the only thing different is the method used to calculate the probabilities. Some of this section will be redundant with the previous note so that each can stand alone.

The fully articulated calculation in Supplementary Note 2 explicitly follows all possible sequences of mergers in the slow Kingman coalescent. But our more efficient calculation here instead leverages two observations:

1. fast-stage outcomes that differ only in which diploid individuals collapse have the same DGS edge-length vector, and
2. in an *m*-lineage Kingman coalescent, the expected edge length ancestral to any specified subset of *j* initial lineages can be computed from the expected total length of all edges subtending those *j* lineages.

Using these, equivalent fast-stage outcomes in the first stage and equivalent descendant subsets in the second stage can be bundled into groups rather than computed individually. This exchangeability keeps us from having to enumerate every possible slow coalescent history, which is a common kind of simplification in SFS-style calculations (Kamm et al., 2017; Wakeley, 2009). We demonstrate each counting step below and recover exactly the same expected edge lengths and DGS probabilities as the full recursion in Supplementary Note 2.

As previously: For a polarized mutation, a DGS bin is written

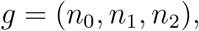

where *n*_0_, *n*_1_, and *n*_2_ are the numbers of homozygous ancestral, heterozygous, and homozygous derived individuals, respectively. We again sample individuals A, B, and C, whose homologs are labeled (0, 1), (2, 3), and (4, 5). We measure edge lengths in coalescent units, and we do not consider anything above the sample MRCA because mutations there would be present in every sampled homolog and produce a monomorphic site.

### C2. Grouping the fast-stage outcomes

At *s* = 0.7, the equilibrium inbreeding coefficient and the corresponding probability of escaping the fast phase without collapse are

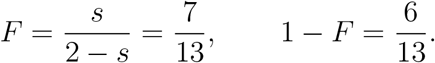

We let *x* be the number of the three sampled diploids whose two homologs collapse during the fast phase. There are

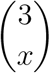

labeled fast-stage outcomes with the same value of *x*. Each possible labeled outcome has probability

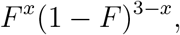

so the total probability of the class containing *x* collapses is just this expression multiplied by the number of possible outcomes that produce *x* collapses, or:

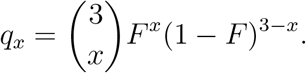

A key observation is that the identities of the collapsed individuals are exchangeable and have no effect on the resulting DGS edge-length vector. So for this example we can reduce the eight labeled fast-stage outcomes in Supplementary Note 2 to just four broader “symmetry classes”:

| $x$ | Possible slow state | Number represented | Class probability $q_x$ | Decimal |
| --- | --- | --- | --- | --- |
| 0 | $\{0, 1, 2, 3, 4, 5\}$ | $\binom{3}{0} = 1$ | 216/2197 | 0.098316 |
| 1 | $\{(01), 2, 3, 4, 5\}$ | $\binom{3}{1} = 3$ | 756/2197 | 0.344105 |
| 2 | $\{(01), (23), 4, 5\}$ | $\binom{3}{2} = 3$ | 882/2197 | 0.401456 |
| 3 | $\{(01), (23), (45)\}$ | $\binom{3}{3} = 1$ | 343/2197 | 0.156122 |

These four class probabilities sum to

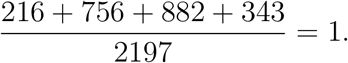

A class with *x* collapsed diploids begins with

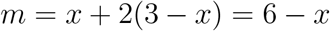

lineages in the slow stage: one lineage from each collapsed diploid and two lineages from each uncollapsed diploid.

### C3. Expected edge length for one specified descendant subset

Suppose that the slow coalescent begins with *m* labeled lineages. Every edge in the resulting genealogy can be classified by the subset of those initial lineages that descend from it.

For example, an edge may be ancestral to one initial lineage, to a particular pair of lineages, or to a particular set of three, etc. A mutation on an edge ancestral to *j* initial slow lineages would be inherited by the sampled homologs descended from those lineages (Wakeley, 2009).

Here we will leverage a well-known edge-length result under the Kingman coalescent. Let 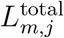 refer to the random combined length of all edges ancestral to exactly *j* of the *m* initial lineages. Using a coalescent rate of 1,

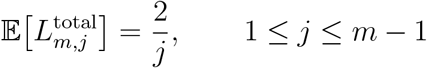

(Dahmer and Kersting, 2015; Fu, 1995; Griffiths and Tavaré, 1998; Wakeley, 2009).

This total 2*/j* represents the expected summed length of all specified subsets of *j* initial slow lineages. But for the DGS calculation, we are interested in the expected length associated with one specified subset. For example, in the slow state

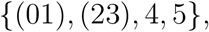

we may want to calculate the expected edge length ancestral specifically to the subset

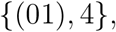

rather than the combined length ancestral to all possible pairs. There are

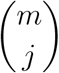

specified subsets containing *j* of the *m* initial lineages. Every subset of a given size has the same expected ancestral edge length, so to get this particular expected length, just divide the expected total length equally among these subsets.

More generally, if we define *w*_*m,j*_ as the expected length of edges ancestral to some specified subset of *j* initial lineages, then

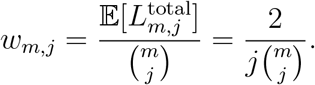

So as an example, consider the two-collapse outcome whose representative slow state is

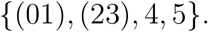

This state begins with *m* = 4 slow lineages. The expected total length of all edges ancestral to exactly two of these lineages is

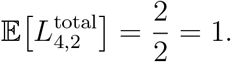

There are

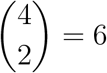

possible specified pairs of these initial slow-phase lineages. Because the six pairs are exchangeable, the expected edge length ancestral to any one specified pair is

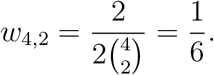

### C4. Counting the possible subsets that produce each DGS bin

We now want to count how many possible descendant subsets produce the same diploid genotype configuration. Consider a fast-stage class with *x* collapsed diploids and

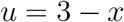

uncollapsed diploids.

A specific descendant subset is some particular collection of those lineages. For example, in the two-collapse outcome

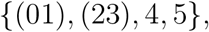

the collections

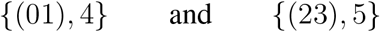

are two different specified subsets, even though mutations on their ancestral edges produce the same DGS configuration (i.e., they are exchangeable in the DGS calculations).

We can group the possible subsets based on how they partition the initial slow-phase lineages. Among the *x* collapsed diploids, let

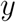

be the number included in the subset. Each contributes one slow lineage to the subset but produces one homozygous-derived individual.

In contrast, each of the *u* uncollapsed diploids contributes two separate slow lineages. Relative to a given subset, an uncollapsed diploid must fall into exactly one of three categories:

*h* (“heterozygote”) exactly one of its two homolog lineages is included,

*d* (“double”) both homolog lineages are included, and

*u* − *h* − *d* (“absent”) neither homolog lineage is included.

The subset contains

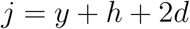

slow lineages. The term *y* counts one slow lineage for each included collapsed diploid, *h* counts one lineage from each singly represented uncollapsed diploid, and 2*d* counts two lineages from each doubly represented uncollapsed diploid.

It is important to recognize at this point that each (*y, h, d*) will map to a single DGS configuration. We can count the subsets corresponding to a given (*y, h, d*) in four steps:

First, choose which *y* of the *x* collapsed diploids have their lineage included:

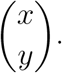

Second, choose which *h* of the *u* uncollapsed diploids contribute exactly one homolog:

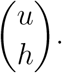

Third, from the remaining *u* −*h* uncollapsed diploids, choose which *d* contribute both homologs:

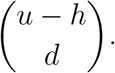

The rest of the *u* −*h* −*d* uncollapsed diploids then do not contribute either homolog, so we don’t have to consider them.

Finally, for each of the *h* diploids contributing exactly one homolog, there are two choices for which homolog is included. These choices are independent, giving

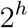

possible choices of homologs.

We can multiply these independent choices to give the number of specified subsets that share the same (*y, h, d*) description:

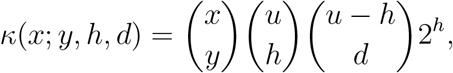

and we can now add this contribution to its specific DGS bin:

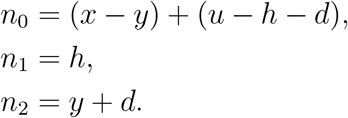

Every *κ*(*x*; *y, h, d*) specified subset contains

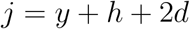

slow lineages. From Section C3, the expected edge length ancestral to any one such specified subset is

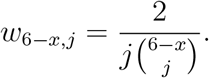

So the combined expected edge length contributed by all subsets with the same corresponding (*y, h, d*) is

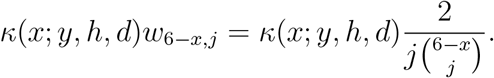

This contribution is assigned to the DGS bin

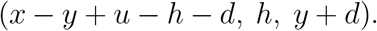

The tables below loop through every possible combination of (*y, h, d*), for each fast-stage class.

### C5. No-collapse class: x = 0, u = 3, m = 6

Here *y* = 0 in every row, since each of the six sampled homologs is initially a separate slow lineage.

| $y$ | $h$ | $d$ | $j$ | $\kappa$ | $w_{6,j}$ | DGS bin | $\kappa w_{6,j}$ |
| --- | --- | --- | --- | --- | --- | --- | --- |
| 0 | 0 | 1 | 2 | 3 | 1/15 | (2, 0, 1) | 1/5 |
| 0 | 0 | 2 | 4 | 3 | 1/30 | (1, 0, 2) | 1/10 |
| 0 | 1 | 0 | 1 | 6 | 1/3 | (2, 1, 0) | 2 |
| 0 | 1 | 1 | 3 | 12 | 1/30 | (1, 1, 1) | 2/5 |
| 0 | 1 | 2 | 5 | 6 | 1/15 | (0, 1, 2) | 2/5 |
| 0 | 2 | 0 | 2 | 12 | 1/15 | (1, 2, 0) | 4/5 |
| 0 | 2 | 1 | 4 | 12 | 1/30 | (0, 2, 1) | 2/5 |
| 0 | 3 | 0 | 3 | 8 | 1/30 | (0, 3, 0) | 4/15 |

To highlight an example, the row (*h, d*) = (1, 1) counts the possible subsets containing a single homolog from one individual, both homologs from one individual, and neither homolog from the remaining individual. There are

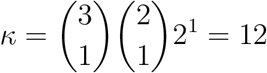

such subsets. Each contains *j* = 1 + 2 = 3 slow lineages and has expected length *w*_6,3_ = 1*/*30, giving 12*/*30 = 2*/*5 in bin (1, 1, 1).

Summing the rows gives the no-collapse column

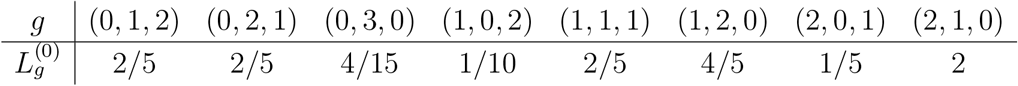

and total length 137*/*30 = 2(1 + 1*/*2 + 1*/*3 + 1*/*4 + 1*/*5), the expected total tree length for six Kingman lineages.

### C6. One-collapse class: x = 1, u = 2, m = 5

Take {(01), 2, 3, 4, 5} as the example one-collapse state. The other two single-collapse states are accounted for by multiplying by 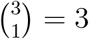 as *q*_1_, since the one-collapse states are exchangeable.

| $y$ | $h$ | $d$ | $j$ | $\kappa$ | $w_{5,j}$ | DGS bin | $\kappa w_{5,j}$ |
| --- | --- | --- | --- | --- | --- | --- | --- |
| 0 | 0 | 1 | 2 | 2 | 1/10 | (2, 0, 1) | 1/5 |
| 0 | 0 | 2 | 4 | 1 | 1/10 | (1, 0, 2) | 1/10 |
| 0 | 1 | 0 | 1 | 4 | 2/5 | (2, 1, 0) | 8/5 |
| 0 | 1 | 1 | 3 | 4 | 1/15 | (1, 1, 1) | 4/15 |
| 0 | 2 | 0 | 2 | 4 | 1/10 | (1, 2, 0) | 2/5 |
| 1 | 0 | 0 | 1 | 1 | 2/5 | (2, 0, 1) | 2/5 |
| 1 | 0 | 1 | 3 | 2 | 1/15 | (1, 0, 2) | 2/15 |
| 1 | 1 | 0 | 2 | 4 | 1/10 | (1, 1, 1) | 2/5 |
| 1 | 1 | 1 | 4 | 4 | 1/10 | (0, 1, 2) | 2/5 |
| 1 | 2 | 0 | 3 | 4 | 1/15 | (0, 2, 1) | 4/15 |

Some bins receive more than one row. For example,

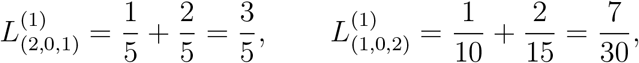

and

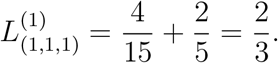

The complete one-collapse vector is

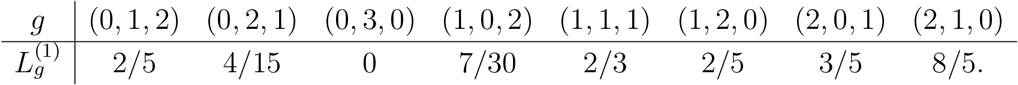

Its entries sum up to 25*/*6 = 2(1 + 1*/*2 + 1*/*3 + 1*/*4), the five-lineage Kingman total.

### C7. Two- and three-collapse classes

For two collapsed diploids, we can consider the example state {(01), (23), 4, 5}. Here *x* = 2, *u* = 1, and *m* = 4.

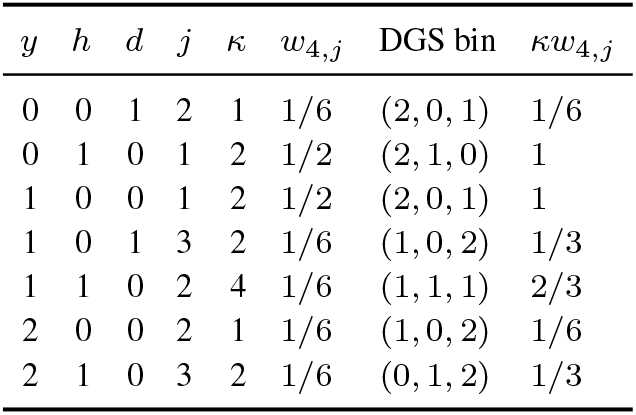

So

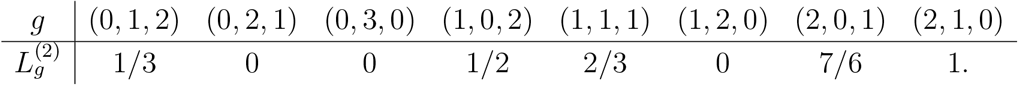

The entries sum to 11*/*3 = 2(1+1*/*2+1*/*3), matching the total length expected under the Kingman coalescent for four lineages and matching the check from the full recursion in Supplementary Note 2.Again exchangeability applies, so this example 2-collapse state will be multiplied by ^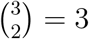^ in a later step (see C9).

When all three diploids collapse, the state is {(01), (23), (45)}, so *x* = 3, *u* = 0, and *m* = 3.

There are only two subset types under this scenario:

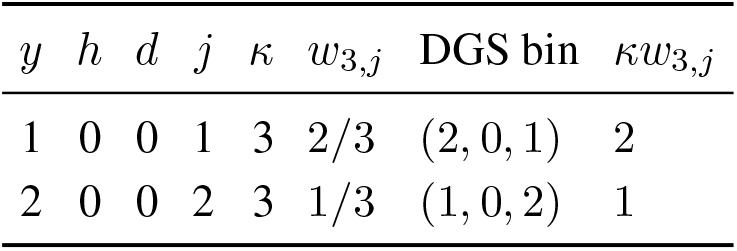

So 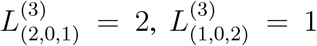, and all other entries are zero. Their sum is 3, the three-lineage Kingman expected total length.

### C8. Conditional edge lengths for all four symmetry classes

Consolidating the row sums above gives us exactly the same conditional edge-length table as the articulated recursion:

| DGS bin | $x = 0$ | $x = 1$ | $x = 2$ | $x = 3$ |
| --- | --- | --- | --- | --- |
| (0, 1, 2) | 2/5 | 2/5 | 1/3 | 0 |
| (0, 2, 1) | 2/5 | 4/15 | 0 | 0 |
| (0, 3, 0) | 4/15 | 0 | 0 | 0 |
| (1, 0, 2) | 1/10 | 7/30 | 1/2 | 1 |
| (1, 1, 1) | 2/5 | 2/3 | 2/3 | 0 |
| (1, 2, 0) | 4/5 | 2/5 | 0 | 0 |
| (2, 0, 1) | 1/5 | 3/5 | 7/6 | 2 |
| (2, 1, 0) | 2 | 8/5 | 1 | 0 |

### C9. Weighting the classes across the fast phase

The expected edge length in bin *g* is now a four-term sum of the single-case calculations above along with their multiplicities due to exchangeability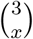 (contained in *q*_*x*_),

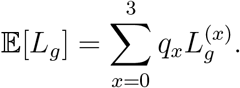

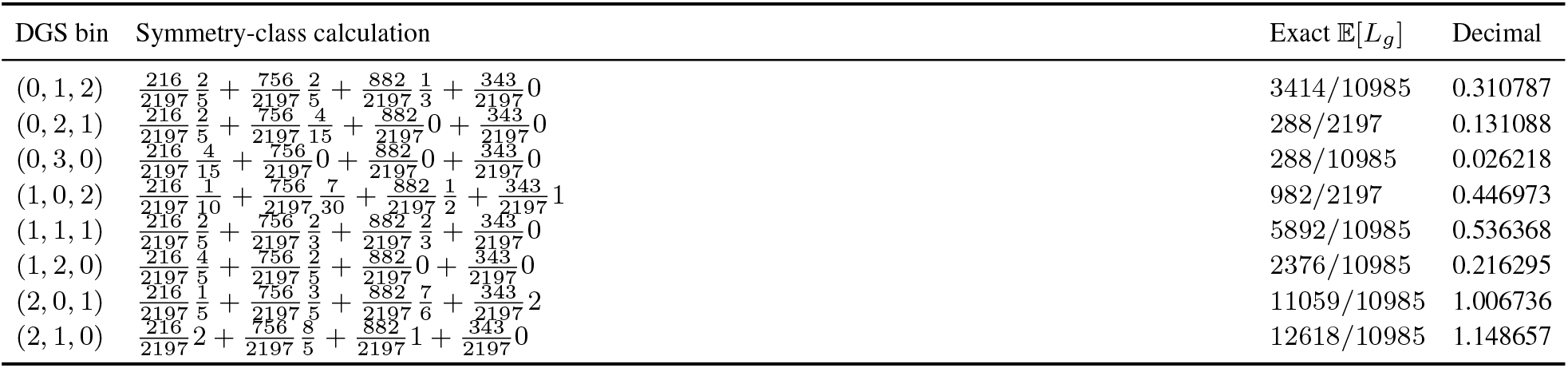

The total weighted polymorphic edge length is again

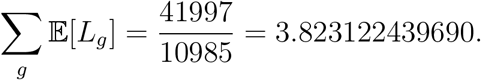

### C10. Final DGS probabilities

Conditional on observing a segregating mutation, the DGS probability is

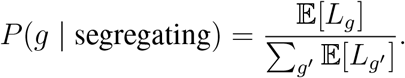

| DGS bin | Weighted edge length | Decimal length | DGS probability |
| --- | --- | --- | --- |
| (0, 1, 2) | 3414/10985 | 0.310787 | 0.081292 |
| (0, 2, 1) | 288/2197 | 0.131088 | 0.034288 |
| (0, 3, 0) | 288/10985 | 0.026218 | 0.006858 |
| (1, 0, 2) | 982/2197 | 0.446973 | 0.116913 |
| (1, 1, 1) | 5892/10985 | 0.536368 | 0.140296 |
| (1, 2, 0) | 2376/10985 | 0.216295 | 0.056575 |
| (2, 0, 1) | 11059/10985 | 1.006736 | 0.263328 |
| (2, 1, 0) | 12618/10985 | 1.148657 | 0.300450 |

These probabilities sum to one and are identical to those from the full recursion in Supplementary Note 2 (see section S8).

### C11. Direct correspondence with the optimized implementation

`selfdgs` follows the arithmetic above directly. For each *x* = 0, …, *n*, it performs the following steps:

1. compute the fast-stage class probability

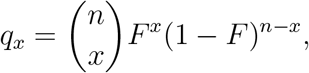
2. set *u* = *n* − *x* and *m* = 2*n* − *x*,
3. loop over *y* = 0, …, *x, h* = 0, …, *u*, and *d* = 0, …, *u* − *h*,
4. set

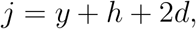

(skip the case of *j* = 0 or *j* = *m*),
5. compute the subset multiplicity

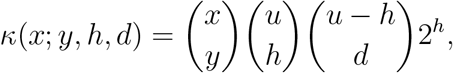
6. compute the expected length for one specified subset,

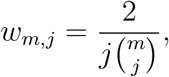
7. add

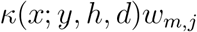

to the class edge-length entry for

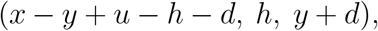

and
8. multiply the completed class edge-length vector by *q*_*x*_ and add it to the expectation across fast-stage classes.

